# Nanopore DNA Sequencing and Genome Assembly on the International Space Station

**DOI:** 10.1101/077651

**Authors:** Sarah L. Castro-Wallace, Charles Y. Chiu, Kristen K. John, Sarah E. Stahl, Kathleen H. Rubins, Alexa B. R. McIntyre, Jason P. Dworkin, Mark L. Lupisella, David J Smith, Douglas J. Botkin, Timothy A. Stephenson, Sissel Juul, Daniel J. Turner, Fernando Izquierdo, Scot Federman, Doug Stryke, Sneha Somasekar, Noah Alexander, Guixia Yu, Christopher E. Mason, Aaron S Burton

## Abstract

The emergence of nanopore-based sequencers greatly expands the reach of sequencing into low-resource field environments, enabling *in situ* molecular analysis. In this work, we evaluated the performance of the MinION DNA sequencer (Oxford Nanopore Technologies) in-flight on the International Space Station (ISS), and benchmarked its performance off-Earth against the MinION, Illumina MiSeq, and PacBio RS II sequencing platforms in terrestrial laboratories. Samples contained mixtures of genomic DNA extracted from lambda bacteriophage, *Escherichia coli* (strain K12) and *Mus musculus* (BALB/c). The in-flight sequencing experiments generated more than 80,000 total reads with mean 2D accuracies of 85 – 90%, mean 1D accuracies of 75 – 80%, and median read lengths of approximately 6,000 bases. We were able to construct directed assemblies of the ~4.7 Mb *E. coli* genome, ~48.5 kb lambda genome, and a representative *M. musculus* sequence (the ~16.3 kb mitochondrial genome), at 100%, 100%, and 96.7% pairwise identity, respectively, and *de novo* assemblies of the lambda and *E. coli* genomes generated solely from nanopore reads yielded 100% and 99.8% genome coverage, respectively, at 100% and 98.5% pairwise identity. Across all surveyed metrics (base quality, throughput, stays/base, skips/base), no observable decrease in MinION performance was observed while sequencing DNA in space. Simulated runs of in-flight nanopore data using an automated bioinformatic pipeline and cloud or laptop based genomic assembly demonstrated the feasibility of real-time sequencing analysis and direct microbial identification in space. Applications of sequencing for space exploration include infectious disease diagnosis, environmental monitoring, evaluating biological responses to spaceflight, and even potentially the detection of extraterrestrial life on other planetary bodies.

## Introduction

The ability to sequence DNA off-planet would address a critical need for ensuring crew health and safety during space exploration missions. Durations for Mars missions are likely to range from 18 to 36 months, with 12 to 18 months of that time spent in transit between the planets, based on current propulsion technologies. Considering the time required to reach Mars and that feasible launch windows for Mars only occur every 26 months due to orbital dynamics, intervention from Earth during the course of a Mars mission will be limited to electronic communication. It is also known, though not well understood, that the human immune response becomes depressed in response to spaceflight (*1*), and gene expression-modulated increases in microbial virulence have been observed (*2*). Thus, there is a clear need for in-flight diagnostic capability to ensure that any infections that arise are treated appropriately, including determining whether or not antibiotics should be administered, and if so, which ones (e.g., anti-fungal, antiviral, anti-microbial). Beyond the transcription level changes observed for microbes, it is also unknown how microbial populations will change over the course of a multi-year mission with increased exposure to ionizing radiation and microgravity during transit, both in terms of population ecology and genetic mutations. What is known, however, is that microbiome changes can have significant effects on human health on Earth. There is also the possibility that, if life ever existed on Mars, it could share a common ancestry with DNA-based life on Earth. If so, sequencing genetic material extracted from Mars would be the surest way to determine whether any DNA detected was contamination from Earth or if it was in fact from indigenous life on Mars.

Sequencing is the only currently existing technology that could be used to address all of these questions (disease diagnosis, population metagenomics, gene expression changes, accumulation of genetic mutations, and characterization of life on another planetary surface), potentially with a single instrument. The commercially available MinION™ DNA sequencer (Oxford Nanopore Technologies, Oxford, UK) is small (9.5 × 3.2 × 1.6 cm) and lightweight (86 grams). Power and data transfer occur through a single USB connection to a computer or tablet, making the MinION exceptionally suited for use in remote locations, including space (*3, 4*). To evaluate the performance of the MinION in a relevant space environment, we tested it aboard the International Space Station (ISS). Orbiting 400 km above the Earth and travelling at 28,000 km/h, the ISS is in constant freefall and maintains a continuous microgravity environment. In addition to being testbed for technology to be used in space, the ISS shares many features with remote environments on Earth, such as limited power capacity, re-supply only through extraordinary and costly efforts, and limitations on equipment and supplies requiring specific characteristics in ruggedness and size. Here we describe the results of nanopore DNA sequencing experiments performed aboard the ISS on samples containing a mixture of genomic DNA extracted from a virus (*Enterobacteria phage lambda*), a bacterium (*Escherichia coli*), and a host organism (mouse). In parallel, we performed control experiments on the ground and made cross-platform comparisons with genomic sequence data obtained from the same samples on Illumina MiSeq (Illumina, San Diego, CA) and PacBio (Pacific Biosciences, Menlo Park, CA) instruments. Simulated analyses of in-flight data using an automated metagenomic pipeline and *de novo* genomic assemblies on the cloud and laptop demonstrate the feasibility of on-board, real-time analysis and assembly for future sequencing applications in space.

## Results

### Flight and Ground Control sequencing with the MinION

We launched the MinION to the ISS to evaluate its functionality in the microgravity environment, under the payload mission name “Biomolecule Sequencer”. Four Biomolecule Sequencer experiments were conducted, on August 26^th^, September 3rd, September 7th, and September 13th, 2016 (Supplementary Table 1). Identical, simultaneous sequencing runs were performed on the ground to mitigate variables not directly related to spaceflight and allow comparisons to the spaceflight-generated sequence data and performance metrics of the MinION device. For each run, a frozen sample containing ready-to-sequence DNA libraries was thawed and loaded on a new MinION flow cell, and the sequencing run was initiated using MinKNOW software (Oxford Nanopore Technologies, Inc.). All samples contained equimolar inputs of lambda, *E. coli* and mouse genomic DNA. A total of six distinct 6-hour sequencing runs (three in-flight and three on the ground) from a single bulk sequencing library preparation, and two 48-hour runs (one in-flight and one on the ground) from a second library preparation were performed. Of the eight runs, all but one (ground control #2) produced good yields of high quality sequencing data. Upon completion of sequencing in-flight, all FAST5/HDF files produced were downloaded from the ISS to Earth. The flight and ground data were then analyzed using a number of open-source and custom-developed bioinformatic workflows (see Materials and Methods).

A key determinant of the success or failure of sequencing on the MinION is the number of active pores identified during the MUX scan performed at the initiation of sequencing. Previous vibration testing on the ground suggested that at least ~70% of nanopores in the R7 flow cells should be active after launch vibration (*5*). Though there was considerable variability in the number of active pores, no statistically significant decrease in the number of active pores in the flow cells used on the ISS was observed compared with the ones used on the ground (Supplementary Table 1). The low number of reads during the Ground Control #2 run was not unexpected considering the low number of available pores (n=215). A total of 89,693 reads (containing > 1 billion events), as determined from the totals displayed in the MinKNOW software at the end of each run, were generated across the four ISS experiments, compared with 51,392 from the ground controls. This suggests that MinION sequencing performance on the ISS was comparable to that on the ground.

Initial quality control of the data was conducted using a custom Metrichor workflow for 1D and 2D basecalling and species identification (Figure S1). We performed additional characterization of the in-flight and ground reads by analyzing the k-mer current plots across the four sets of runs (Figure 1), as well as measurements of signal parsing quality with the skips/base (Figure S2) and stays/base (Figure S2). With the exception of ground control #2, the k-mer current profiles, skips/base, and stays/base were within the range of expected values for MinION sequencing and permitted adequate base-calling. We next analyzed the 1D and 2D accuracies of the in-flight and ground reads, along with organisms’ identification from the read data (Figure 2). In general, read accuracies across the ISS and ground runs were comparable, spanning a median of 74-80% (ISS) and 71-78% (ground) median base accuracy for the 1D reads and 83-92% (ISS) and 80-90% (ground) median base accuracy for the 2D reads, though in most cases the accuracies of the in-flight reads were slightly higher. The distributions of reads in both the flight and ground data were found to be fairly evenly distributed across the E. coli genome (Supplementary Figures 3 and 4); median read lengths were typically 5,000 – 6,000 bases (Supplementary Figure 5).

**Figure 1:**
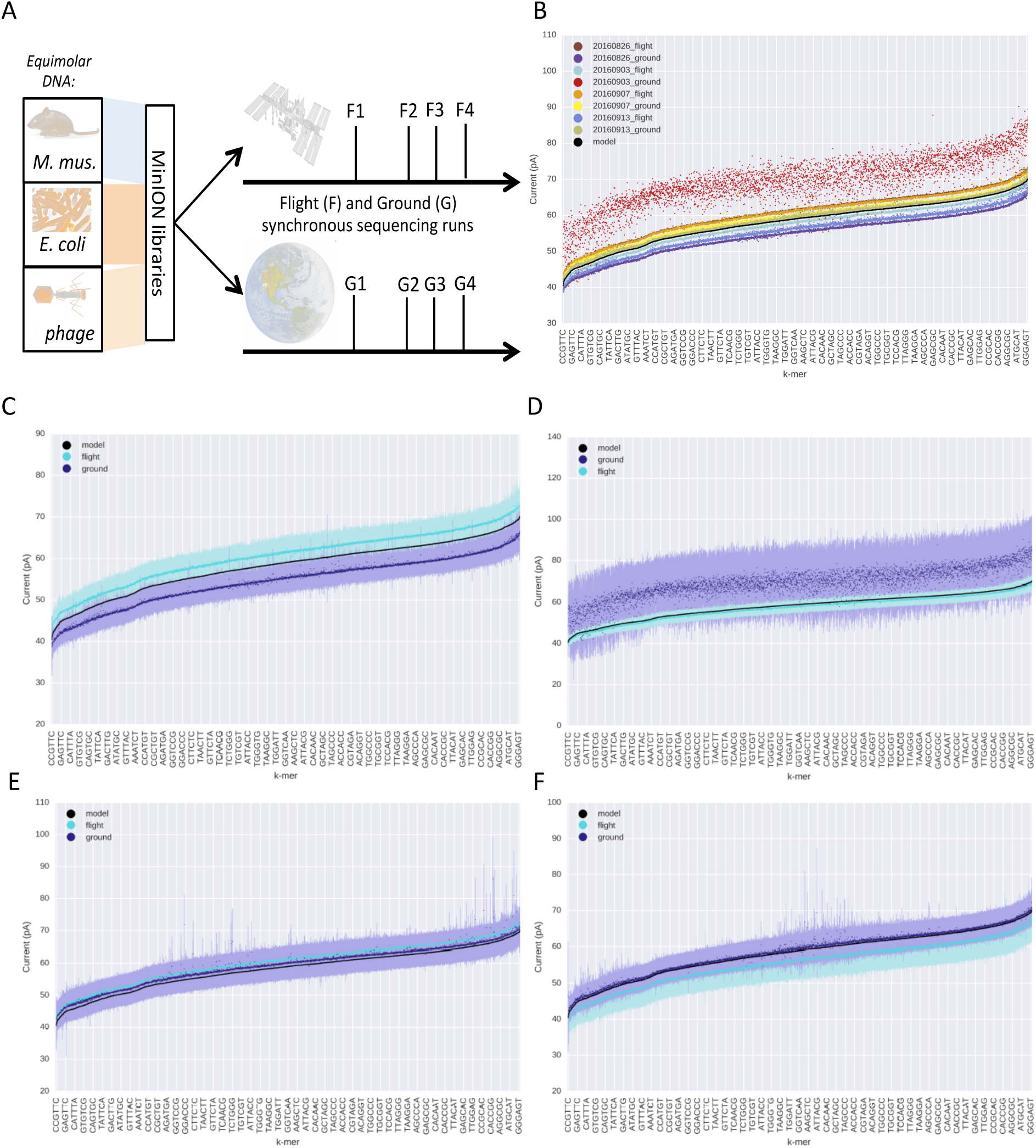
Experimental Setup and electrical current distributions. Mean and standard deviations of currents measured for each k-mer in each of the four runs (top: 1-2, bottom: 3-4), compared to the model from ONT. A subset of the ordered k-mers are listed on the x-axis, although all 4^6 are represented by points. Current distributions all. Mean currents for each k-mer for each of the flight and ground runs (8 total).

**Figure 2:**
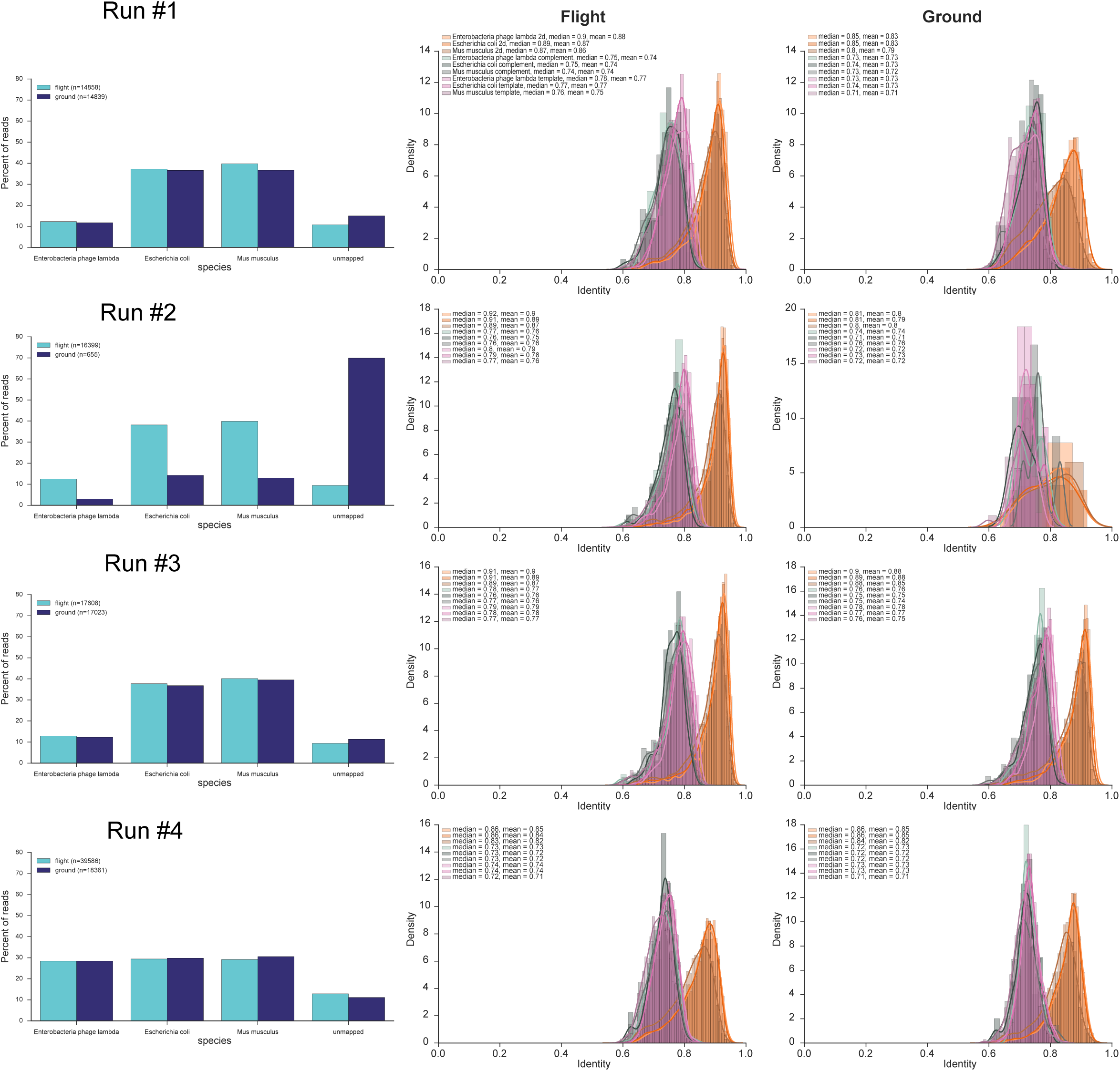
Alignment statistics and base calling for ISS and Ground runs. Results of GraphMap alignments for the four runs, comparing species counts for each species in the mixture (left column), and the fraction identity of aligned segments for the reads generated on the ISS (middle) and on the ground (right) divided by read type and species match. Legend: the 2D reads are shown for mouse, E. coli, and lambda, followed by the 1D reads of the template strand and the complement.

### Benchmarking of the MinION sequencing data against MiSeq and PacBio RSII data

To assess the accuracy of the ISS / ground nanopore MinION data and to establish “gold standard” genomic references for the lambda and *E. coli* genomes, we used the same stock of DNA for replicate sequencing experiments on PacBio RSII and the Illumina MiSeq instruments. We ran 22 SMRT cells across independently sequenced lambda, *E.coli*, and mouse genomic DNA samples on the RSII (1.1M reads with 11,366bp average length) and a full flow cell on the MiSeq (21,214,638 single-end 160 base pair reads). The MiSeq run consisted of separately indexed samples corresponding to genomic DNA from lambda (792, 838 reads), *E. coli* (3,189,286 reads), mouse (7,197,511 reads), and an equimolar mixture of genomic DNA from all 3 organisms (10,035,003 reads) replicating the nanopore-sequenced samples. *De novo* assembly of the *E. coli* genome using the Hierarchical Genome Assembly Process (HGAP, v2) from PacBio reads at 162.7X coverage generated a single contiguous sequence (contig) of size 4,734,145 bp (Figure 3A, Figure S6). A *de novo* assembly from all identified *E. coli* reads on the Illumina MiSeq run produced 245 contigs with coverage of 80.1% of the genome at 99.7% identity to the PacBio assembly (Figure 3B), while directed assembly of *E. coli* Illumina reads to the most closely matched genome in the NCBI nt reference database (*E. coli* K-12, CP014348) resulted in 99.0% coverage at 99.97% identity (Figure 3C). Similarly, genomic assemblies of the lambda genome from PacBio and Illumina data showed 100% concordance. Thus, two orthogonal sequencing methods were used to establish the *E. coli* and lambda genomic references for assessing the accuracy of in-flight and ground nanopore data. A direct alignment of the 2D reads to the *de novo* assembled genomes showed a median 89% and 86% agreement on base calling for the in-flight and ground data, respectively. Overall, we found that the longest molecule from the MinION matched well (90%) to the MiSeq and RSII data, showing that the long, phased genetic data from the MinION is likely amenable to use in analysis of the sequences generated on the ISS.

**Figure 3.**
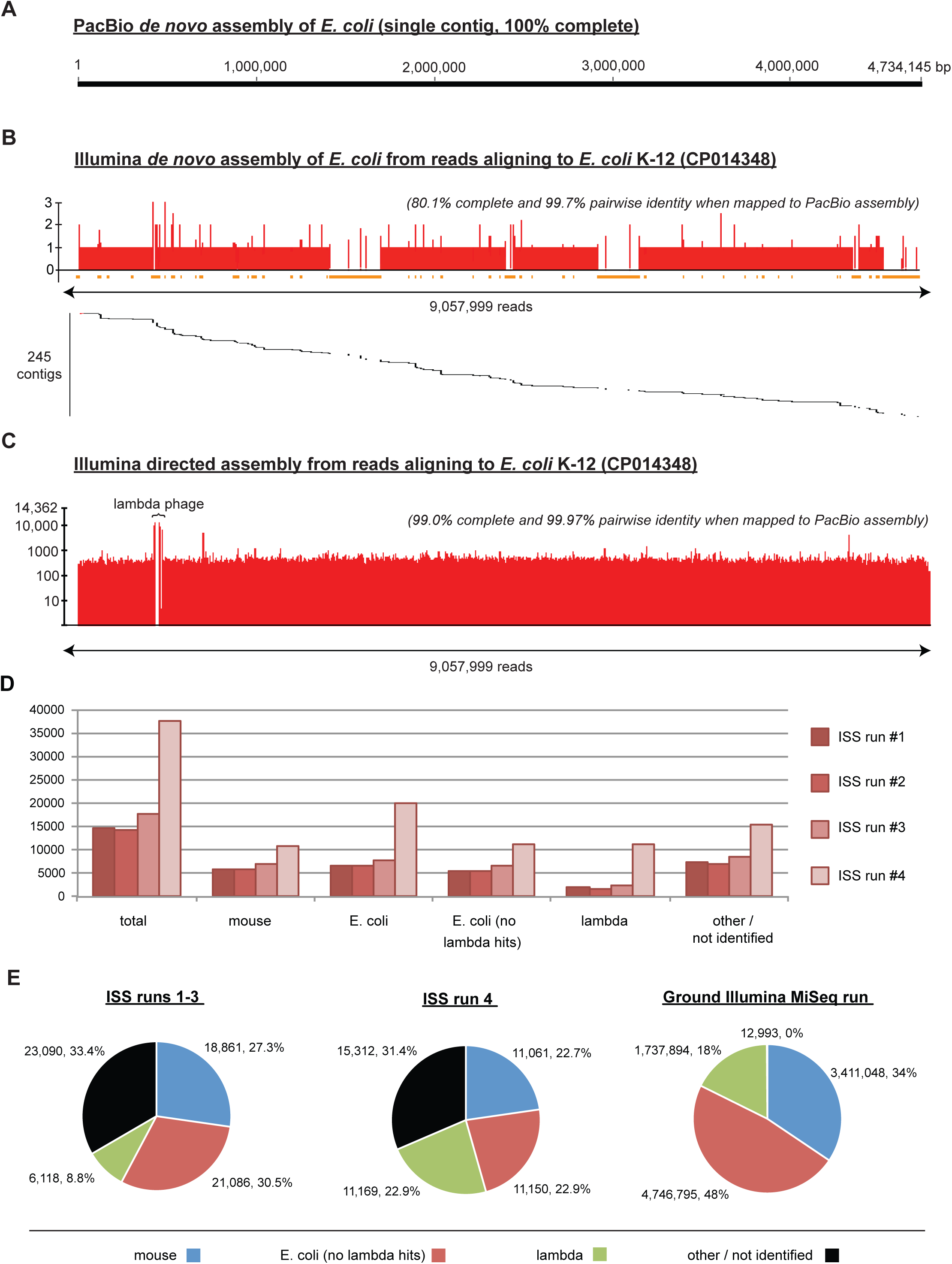
*De novo* genome assembly and cross-platform validation of the ISS nanopore run data. (A) *De novo* assembly of the *E. coli* genome from PacBio reads generates a single full-length contiguous sequence (contig) of length 4,734,145 base pairs (bp). (B) *De novo* assembly of the *E. coli* genome from ~9 million Illumina reads results in 245 mapped contigs (black segments) that assemble into a low-coverage, 80.1% complete genome (red bars) with 99.7% pairwise identity to the “gold standard” PacBio genome assembly. The orange bars denote regions of the genome with no coverage from an Illumina contig. **(C)** Direct assembly of the Illumina reads to the closest matched reference in NCBI GenBank (*E. coli* K-12, CP014348) enables 99.0% genome recovery at 99.97% identity. As the *E. coli* CP014348 reference does not contain integrated lambda prophage, a narrow gap in coverage is observed corresponding to the lambda phage sequence inserted in the PacBio assembly (“lambda phage”). **(D)** Read counts from ISS runs 1-4 corresponding to total, mouse, *E. coli, E. coli* after removal of lambda sequences (“no lambda”), lambda, and other. Note that the distribution of read counts for runs 1 through 3 is quite uniform. **(E)** Pie charts of the distribution of reads in ISS run 1-3 and ISS run 4 in comparison to that obtained from ground Illumina MiSq run of the same sample mixture.

### Accuracy and distribution of in-flight nanopore sequencing reads

The performance of ISS nanopore sequencing runs 1 through 3 were similar, with total yields of 15,000 to 18,000 reads and a relatively lower proportion of lambda reads relative to *E. coli* and mouse reads (Figure 3D and 3E). Unlike the earlier runs, which were sequenced for 6 hours, in-flight run 4 was sequenced for 48 hours, more than doubling the number of reads. Interestingly, the percentage of lambda reads generated from ISS run 4 was 22.9% versus an average of 8.8% in the first 3 runs, and the relative proportions of lambda, mouse, *E. coli* reads were very close to the predicted proportions of 33%/33%/33% that would be expected from sequencing of an equimolar mixture of genomic DNA. For all four runs, approximately one-third of reads were not identified as either lambda, *E. coli*, or mouse, likely due to single-read error rates of 8-20% on the R7 version of flow cells we used (Figure 2). To gauge the improvement of the latest chemistry (R9) for the MinION, we ran an additional equimolar mixture of the three samples on Earth, and we found that the error rates of the latest chemistry, as expected, were indeed lower (5-10%). In contrast, 99.9% of the Illumina reads were properly assigned to one of the 3 organisms (Figure 3E, right), although the relative proportions were less uniform than for flight run #4.

### Metagenomic analysis of in-flight nanopore data using the automated SURPIrt pipeline

To demonstrate the feasibility of analyzing sequencing data on the ISS, we ran simulations of real-time metagenomic analysis from in-flight nanopore data using the automated SURPI*rt* pipeline (Figure 4A). Developed at University of California, San Francisco (UCSF) as a hybrid of two previously published software pipelines, SURPI (*6*) and Metapore (*7*), SURPI*rt* is a bioinformatic pipeline for real-time pathogen detection and genomic characterization from metagenomic nanopore sequencing data. Species-specific reads corresponding to *M. musculus, E. coli*, and lambda from the sample mixtures were initially identified in SURPI*rt* within 1 minute of beginning sequence analysis by MegaBLAST alignment (word size = 16; e-value=1×10^-5). The distribution of detected reads were visualized in real-time as donut charts on a web browser refreshed every 30 s (Figure 4B; Supplementary Video 1).

**Figure 4.**
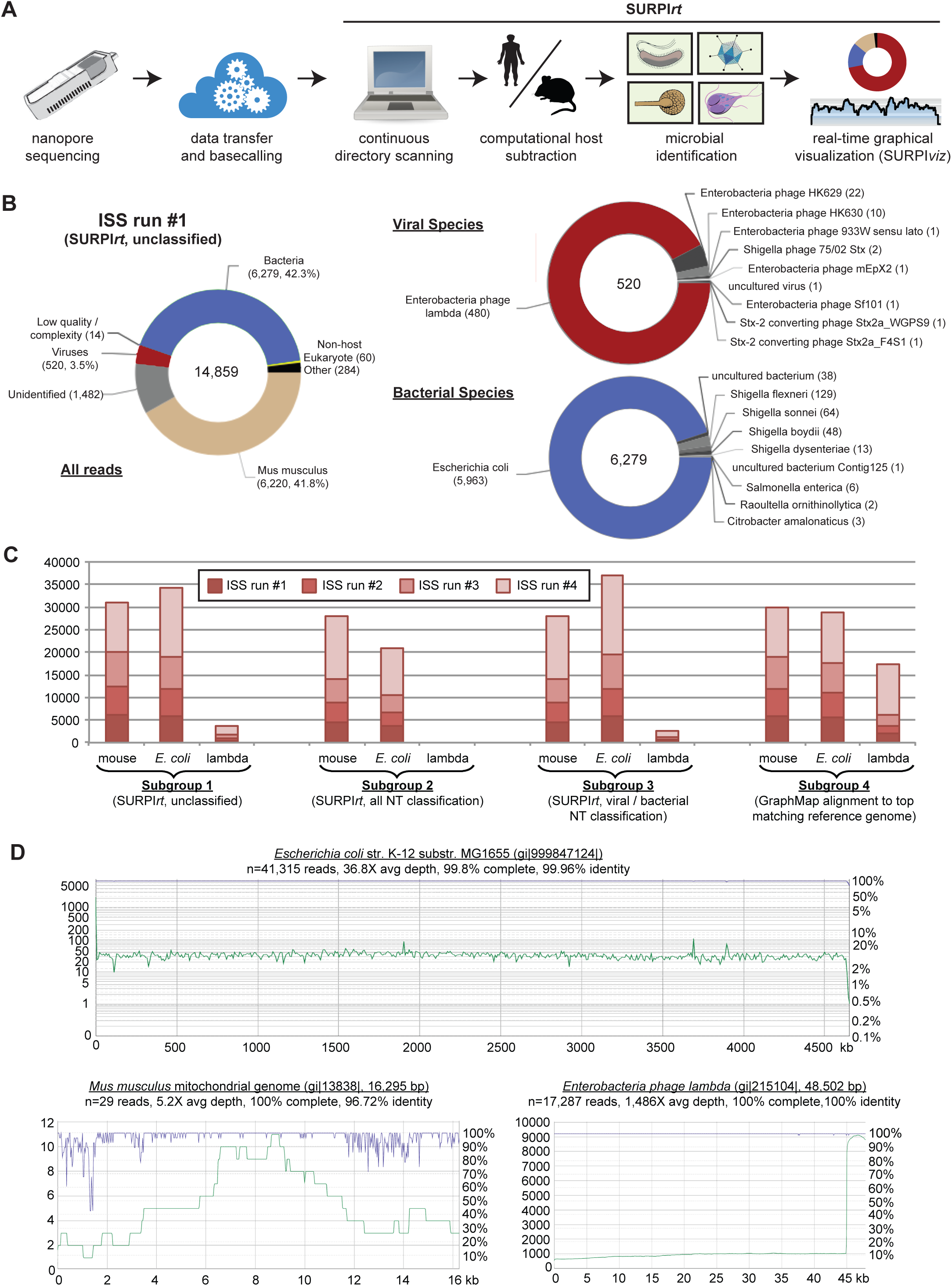
Automated metagenomic analysis of ISS nanopore data. **(A)** Flow chart of the SURPI*rti* bioinformatics pipeline for real-time microbial detection from nanopore data. **(B)** Donut charts of read distributions corresponding to all reads (left), viruses, (upper right), and bacteria (upper right) from ISS run 1. These charts were generated dynamically in real-time as part of a real-time sequencing analysis simulation using SURPI*rt*. **(C)** Stacked distributions of reads from ISS runs 1 through 4 aligning to mouse, *E. coli*, and lambda. Subgroup 1 shows raw SURPI*rt* output in the absence of taxonomic classification, while subgroups 2 and 3 show the effects of classification using the GenBank NT database and separate viral or bacteria databases, respectively. The relative proportions of read counts from SURPI*rt* differ from those obtained by GraphMap alignment to the most closely matched reference genome in NCBI NT (subgroup 4). **(D)** Coverage (green) and pairwise identity plots (purple) of raw nanopore reads mapped the *E. coli* (upper panel), the *M. musculus* mitochondrial (lower left panel), and lambda genomes (lower left panel). Reads are mapped to the most closely matched reference genome identified by SURPI*rt*.

Despite the average error rates of the individual nanopore reads, the vast majority of reads (>95%) were correctly identified to the species level (Figure 4B). Taxonomic classification using a lowest common ancestor (LCA) algorithm (a configurable parameter in SURPI*rt*), while improving specificity, resulted in fewer identified species-specific reads overall, and no lambda reads were detected (Figure 4C, subgroups 1 and 2). The latter finding was attributed to the presence of a lambda prophage insertion in some *E. coli* reference genomes; separate classification of viral and bacterial taxa restored detection of a fraction of the missing lambda reads (Figure 4C, subgroup 3).

Strikingly, the relative proportion of identified lambda reads in the nanopore data was strikingly much less than that observed in the mouse or *E. coli* data (Figure 4C, subgroups 1-3), despite preparation of the sample DNA genomes in equimolar concentrations. This finding was attributed to the presence of the integrated lambda prophage, as long nanopore reads containing lambda insertion sites would be preferentially assigned to *E. coli* rather than lambda. A more uniform distribution of mouse, *E. coli*, and lambda reads was observed when nanopore reads were aligned to the closests matched target genome identified by SURPI*rt* using GraphMap, after ensuring that only single reads were mapped and that lambda sequences were excluded from the set of reads aligning to *E. coli* (Figure 4C, subgroup 4).

SURPI*viz* is a sequence alignment visualization software package designed to receive MegaBLAST and/or GraphMap alignments from SURPI*rt* and display the results in a user-friendly, web-accessible interface. SURPI*viz* results are generated and processed in an automated fashion, and can be viewed in a variety of formats, including heat maps, Krona plots (*8*), coverage maps, and pairwise identity plots (Supplementary Video 2). GraphMap alignment and mapping of raw nanopore reads from flight runs 1 through 4 demonstrated that the 4.5 Mb *E. coli* genome, 47 Kb lambda genome, and 16 Kb *M. musculus* mitochondrial genome (a representative mouse sequence) could be recovered at 96.7-100% pairwise identity by directed assembly to the closest matched reference identified by SURPI*rt* (Figure 5D).

**Figure 5.**
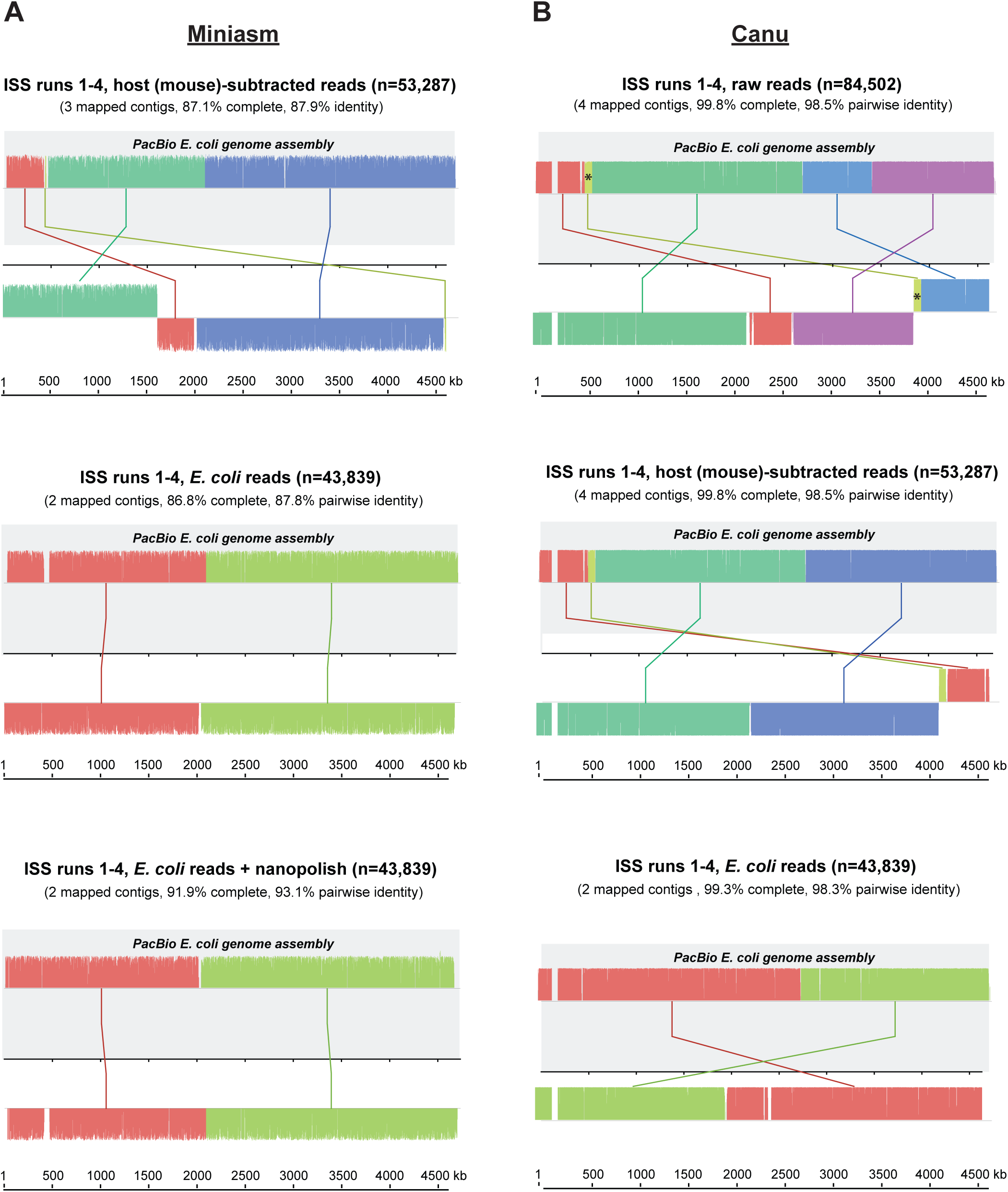
*De novo* assembly of *E. coli* and lambda genomes from ISS nanopore data. Shown using Mauve software are alignments of *de novo* assembled contigs to the PacBio genome assembly used as a “gold standard” reference (gray background). **(A)** Contigs are *de novo* assembled using Miniasm from host (mouse)-subtracted reads (top panel), *E. coli* reads (middle panel), or *E. coli* reads with nanopolish (bottom panel). **(B)** Contigs are *de novo* assembled using Canu from raw reads (top panel), host (mouse)-subtracted reads (middle panel), or *E. coli* reads only (bottom panel). Homologous segments are shown as colored blocks, with blocks that are shifted downward representing segments that are inverted relative to the PacBio reference genome. Similarly colored lines connecting the blocks are used to indicate mapped positions in the reference genome. The asterisk denotes a misassembled region in one of the contigs assembled from a dataset consisting of raw nanopore reads (B, top panel).

### *De novo* genome assembly across the sequencing platforms

To determine if the in-flight nanopore data generated on the ISS could be used for successful *de novo* assembly of the *E. coli* genome, we independently tested two state-of-the-art genome assemblers from nanopore reads, Miniasm (*9*) and Canu (https://github.com/marbl/canu). For the Miniasm assembly, we first filtered out the host reads (mouse) from the ISS data, to leave only those from *E. coli* and lambda. We then used the Miniasm genome assembler followed by nanopolish (https://github.com/jts/nanopolish) to assemble and correct the genome, creating a total of two contigs: 1,636,060bp and 3,042,754 bp. These two contigs add to a total of 4.697 Mb out of an expected size of 4.734 Mb based on the PacBio assembly (98.8% complete), with 97.33% accuracy relative to the PacBio reference genome.

Next, we ran Canu on both a computational server and a laptop to perform successive *de novo* assemblies from nanopore sequencing data corresponding to (i) in-flight runs 1-4, (ii) in-flight runs 1-4, (iii) in-flight runs 1-4 after computational subtraction of mammalian “host” reads corresponding to mouse genome, and (iv) in-flight runs after filtering for reads matching bacteria (Figure 5B). Overall, assemblies were accurate, >99% complete with >98.3% pairwise identity relative to the PacBio reference genome. Although Miniasm had significantly faster run time than Canu (<1 minute versus 2 hours on a 64-core server), Canu appeared to be more accurate, with more complete genomes and improved accuracy relative to Miniasm. *De novo* assembly of nanopore reads using Canu resulted in 100% coverage at 98.5% pairwise identity, an impressive performance despite the presence of a small misassembly in one of the four contigs.

We further tested the ability to run these assemblies not only on computational servers but also on the cloud (the Amazon Elastic Compute Cloud / EC2 platform) and on a laptop. We observed that an 8-core, 32 Gb EC2 instance was sufficient to complete an entire Miniasm assembly within 15 seconds, and a similarly configured laptop with 8 hyperthreaded cores and 64 Gb RAM took 40 sec (Supplementary Figure 7). These results on the cloud and laptop show that ISS, spacecraft, and satellite-based, Earth-free genome sequencing and assembly are possible. As the lambda prophage is inserted into the *E. coli* genome, these data also constitute the first *de novo* assemblies of a complete bacterial and viral genome with 100% accuracy, and indeed, the first genome assemblies of any organism from sequence data generated solely off of planet Earth.

## Discussion

Our results of the first-ever DNA sequencing in space indicate that the performance of the MinION sequencing platform was not adversely affected by transport to the ISS nor by loading or operation in its microgravity environment. The data obtained from sequencing aboard the ISS can readily recapitulate the measurements of nucleic acids from phages (lambda), bacteria (*E. coli*), and mammalian (mouse) DNA on Earth. Indeed, across all three species, the base quality and 2D read alignments were routinely above 85%, with equal or superior performance to the identical replicate libraries and flow cells tested on Earth. These results were also true when comparing the skips/base, stays/base, read length, and GC-content of the data. These sequence reads were also validated against sequencing data obtained on PacBio RSII and Illumina MiSeq instruments to confirm their accuracy and species identity.

Moreover, when using the data produced only from space for a *de novo* assembly, we observed that the *E. coli* reads were sufficient to generate a high-quality genome assembly (2 contigs, >98% identity) from both the Canu and Miniasm+nanopolish assemblers. Excitingly, these data demonstrate that a *de novo* assembly of microbial genomes from raw, unfiltered data containing interfering host background sequences (e.g. mouse genome) is possible. We tested assembler with the lowest computational requirements (Miniasm alone) on both a cloud-based platform (EC2) and also on a laptop. As with sequence data from any platform, the success of unfiltered assembly from nanopore reads will likely depend on the complexity of the metagenomic background in the sample, depth of sequencing, and sequencing error rates, which have been steadily decreasing over time for nanopores (*10*). In aggregate, these current results clearly validate the future implementation of the MinION for rapid, *in situ* diagnostics and microbial identification on the ISS.

As exploration progresses beyond low Earth orbit toward extended missions in cis-lunar space and eventually to Mars, changes and advances in nanopore-based sequencing will be needed. As communication delays increase and data transfer rates decrease, local analysis of sequencing data will be critical. Aspects of this challenge are manifested on Earth in remote locations and point-of-care settings of clinical and public health significance, such as “hot spots” from outbreaks due to Ebola or Zika virus (*11*). In the current study, we used the SURPI*rt* platform to simulate an automated metagenomic analysis of nanopore data in real-time from read processing to microbial identification to genome assembly, and we also showed an entirely cloud-based, rapid (15sec) assembly was possible, highlighting the ability to use these tools and techniques for future missions.

The largest outstanding question for use of the MinION in deep space exploration is flow cell stability over the course of an 18 to 36 month mission. Factors such as extreme temperatures and increased galactic and cosmic radiation exposure are less of a concern for flow cell stability during human missions as crew members will need to be shielded from these conditions as well. For robotic missions, however, enhancements a greater degree of ruggedization will be needed. Stability of nanopore sequencers could be enhanced with the development of more robust membranes in which the pores are embedded or with improvements in the resolving power of solid state nanopores. Assuming the stability challenges are adequately addressed, nucleic acid sequencing would play an important role on crewed missions to Mars to ensure crew health. Once on Mars or on another planetary surface, the sequencing platform would then become a powerful tool for exploration. DNA- or RNA-based life could be rapidly detected and identified, enabling definitive differentiation between Earth-derived contamination and indigenous Martian life. Nanopore sequencing that involves the direct analysis of DNA, rather than indirect products of sequencing-by-synthesis, has been used to detect base modifications in DNA (*12, 13*), and also to sequence RNA (*14*). Direct nanopore analysis has even been used for proteins (*15*). The ability of nanopore analyzers to accommodate a range of polymers increases the chance of detecting extraterrestrial life, which could use different bases or sugars in its genetic material beyond canonical AGCT/U DNA and RNA, greatly increases the value they offer for life detection technology.

In the more immediate future, the MinION holds the potential to greatly improve the rate at which ISS research can be performed by allowing researchers rapid access to data obtained in-flight, rather than having to wait for sample return. With robust experiment planning and some foresight, research projects that required multiple flights over several years could now be performed in a matter of months, as researchers could monitor experiment progress and make adjustments as needed (i.e., cadence of timepoints, identifying a subset of samples that should be returned to Earth for further analysis etc.). Studies of gene expression in-flight would also be enabled by a sequencing platform on the ISS, and could be performed more robustly and with less risk of experiment failure by reducing the need for storage of labile RNA in a freezer, eliminating time constraints posed by organism re-acclimation upon return to Earth, and facilitating more optimal and less arbitrary selection of time-points for sample collection and analysis.

## Conclusion

The observation that the MinION sequencing platform performed equally well (or better) on orbit than it did on the ground without any modifications for spaceflight highlights the potential for nanopore-based sequencing in future space exploration missions. Possible applications of nanopore sequencing in space missions include environmental monitoring, temporal analysis of the ISS microbiome; diagnosis of infectious diseases, personalized medicine and response to countermeasures, tracking of genetic, epigenetic and gene expression changes in response to spaceflight, assessment of Earth-derived biological contaminants on planetary surfaces, and even *in situ* characterization of indigenous DNA or RNA-based life beyond Earth. The validation of the MinION in the spaceflight environment is a critical first step for bringing molecular biology to the space age.

## Materials and Methods

### Spaceflight Hardware

The full payload system included the following items: two MinION devices, a USB 3.0 cable, nine R7.3 flow cells (Oxford Nanopore Technologies), nine sample syringes containing ground-prepared genomic DNA, nine empty sample syringes for air bubble removal, and a configuration flow cell. A pipette kit including 10, 100, and 1,000 μl Rainin positive displacement pipettes (Mettler Toledo, Oakland, CA) and associated tips was included for contingency purposes if the syringe was not sufficient for bubble removal. These items were all launched from Cape Canaveral Air Force Base on the SpaceX CRS-9 Dragon capsule on July 18, 2016. The MinKNOW™ software (Oxford Nanopore Technologies) required for operation of the MinION was loaded on a Microsoft Surface Pro 3 tablet and delivered to the ISS on the Orbital ATK OA-6 Cygnus Space Station Resupply Vehicle, which launched on March 22, 2016.

### Library Preparation and Sequencing

Samples analyzed in this study contained sequence libraries prepared from mixtures of female mouse BALB/C (Zyagen, San Diego, CA), *Escherichia coli* K-12 (Zyagen), and *N*^6^-methyladenine-free bacteriophage lambda (New England Biolabs, Ipswich, MA) genomic DNA. These species were chosen based on their being model mammal, bacterium, and virus, respectively. The samples were aliquoted for library preparation on three platforms: Oxford Nanopore Technologies (ONT) MinION (v6), Illumina MiSeq (v2), and the Pacific Biosciences RSII.

### MinION Library Preparation and Sequencing (Ground and ISS)

Aliquots of DNA for mouse, *E. coli*, and lambda libraries were sheared individually using Covaris g-TUBEs (Covaris, Boston, MA) by centrifugation at 4,800 × g for 2 min to produce fragments that were predominantly 8 kb. Fragmentation was verified using a 2100 Agilent Bioanalyzer (Agilent Technologies, Santa Clara, CA). Sheared mouse, *E. coli,* and lambda DNA samples were quantified using a Qubit 2.0 fluorometer (Thermo Fisher Scientific, Waltham, MA) and combined in equal abundances, targeting 1.5 µg total (0.5 µg each). Mixed DNA samples then underwent treatment to repair residual nicks, gaps, and blocked 3’ ends using Formalin-Fixed, Paraffin-Embedded (FFPE) DNA Repair Mix (New England Biolabs), with modifications to the manufacturer’s protocol to include a 0.5X Agencourt AMPure XP system magnetic bead clean-up (Beckman Coulter Genomics, Brea, CA) to remove small fragments of DNA. The repaired DNA was then prepared for sequencing according to Oxford Nanopore Technologies’ procedure for the SQK-MAP-006 Kit. Following library preparation, individual samples were mixed with a 102 ng aliquot of pre-sequencing library (150 µl) with 162.5 µl of 2X running buffer, 6.5 µl fuel mix (Oxford Nanopore Technologies) and 156 µl molecular-grade sterile deionized water to a total volume of 450 µl. This volume was loaded into a 1 ml syringe and the syringe was capped. A total of 18 samples were prepared: nine flight samples and nine ground control samples. Capped syringes containing the samples were packaged with an identical 1 ml syringe for potential bubble removal and syringe tips were placed inside of a large plastic tube to facilitate transport. After syringe loading, the tubes were stored at −80 °C.

### Ground Processing of Cold Stowage Hardware

The nine flight samples were removed from the −80 °C freezer, placed on dry ice in a Styrofoam cooler, and shipped overnight to the launch processing facility at Kennedy Space Center (KSC). Once at KSC, the samples were maintained in a −80 °C freezer until transfer to the powered freezer on the SpaceX Dragon Capsule. To more closely approximate the handling of the flight samples, the nine ground control samples were also removed from the freezer, placed on dry ice in a Styrofoam cooler and allowed to sit on the laboratory bench for the same amount the time that the flight samples were out of the freezer during transit. Eighteen total R7.3 flow cells from the same manufacturing lot were shipped directly from the manufacturer to KSC for flight (9) and JSC to serve as ground controls (9). Upon receipt, the flow cells were stored at +4 °C until the time of launch. The flow cells and samples were maintained within the launch processing labs at KSC until 48 h before the launch, at which point they were loaded onto the vehicle.

### Launch and ISS Stowage Conditions

Flow cells were launched in a double cold bag in which the temperature was maintained between + 2 and + 8 °C. The DNA samples were launched in a powered freezer in which temperatures were maintained between −80 and −90 °C. Upon docking with the ISS, the flow cells and DNA samples were transferred to refrigerator (+2 to +8 °C) and freezer (−80 to −90 °C) dewars within the Minus Eighty Degree Laboratory Freezer for ISS (MELFI), respectively. All other items were stored at ambient ISS conditions.

### Sequencing Experiments

Following complete charging of the Surface Pro 3, it was connected to the MinION via USB cable. Prior to the first sequencing experiment, a configuration test cell was inserted into the MinION device for the configuration test MAP_CTC_Run.py in the MinKNOW software version 0.51.1.39 b201511042121 to ensure proper data exchange between the MinION device and the Surface Pro 3. This version of MinKNOW software was modified so that internet connectivity was not required.

To initiate the experiment, a flow cell and DNA sample syringe tube were collected from their respective MELFI dewars and allowed to equilibrate to ambient temperature (10 - 60 min). The flow cell was inserted into the sequencing device and the bubble removal syringe was used to remove the air immediately adjacent to the sample loading port (~15 μl of air). Flow cell priming was performed by loading approximately 250 μl of the sample containing fuel mix and running buffer and allowing 10 min to elapse prior to loading the remaining 250 μl of sample. Sequencing was initiated using MAP_Lambda_Burn_In_Run_SQK_MAP006.py protocol selected from the MinKNOW software. Real-time communications with the ISS allowed the ground control flow cells and samples to be retrieved, temperatures equilibrated, and injected for priming and final sample loading in a synchronous manner.

### PacBio RSII Library Preparation and Sequencing

Single Molecule, Real-Time (SMRT) sequencing libraries were prepared using the SMRTbell Template Prep Kit 1.0 (Pacific Biosciences) and 20 kb Template Preparation Using BluePippin Size-Selection System protocol (Pacific Biosciences). 5 μg of each sample were used. Library quality and quantity were determined using an Agilent 2200 TapeStation and Qubit dsDNA BR Assay (Life Technologies), respectively. Sequencing was conducted using P6-C4 chemistry and a v3 SMRT Cell (Pacific Biosciences) at Weill Cornell Medicine.

### Illumina MiSeq Library Preparation and Sequencing

Sequencing libraries were prepared from 1 ng of sample using the NexteraXT kit (Illumina) according to the manufacturer’s protocol. Libraries were indexed with dual 8-nt barcodes on each end of the sequencing amplicon. In total, 4 dual-indexed DNA sequencing libraries were constructed, corresponding to lambda bacteriophage, *E. coli*, mouse, and an equimolar mixture of DNA from the 3 organisms. Libraries were quantitated using the Agilent Bioanalyzer and Qubit spectrophotometer and sequenced on an Illumina MiSeq as a 1 × 160 base pair (bp) single-end sequencing run. The approximate percentage of the run allocated to each library as determined by the quantified input concentration was 3% for lambda bacteriophage, 17% for *E. coli*, 30% for mouse, and 50% for the equimolar mixture.

### Data Analysis

#### Basecalling

Basecalling was performed using the Metrichor workflow “2D Basecalling for SQK-MAP006 - v1.107”.

#### Read Selection

We used a custom shell script to extract one read from each fast5 file for further quantification and analysis, selecting the 2D read where available, and the higher quality of the 1D template or complement read where not.

### GraphMap and Calculation of Species Counts/Proportions

As described in previously (*5*), we aligned to a combined *Enterobacteria lambda phage* (NCBI reference sequence NC_001416.1), *Escherichia coli* (NCBI reference sequence NC_000913.3), and *Mus musculus* (mm10, GRCm38.p4) genome using GraphMap version 0.3.0, with the command “graphmap align -r $ref -d $fi -o $name.sam”, which saves the top result for each read. We used the results to count the number of reads mapping to each of the three species and the fraction identity between reads and references.

For comparison of the relative species proportions in the sample mixture between the nanopore in-flight runs and the Illumina data, we separately aligned the Illumina and nanopore reads to the *Mus musculus, E. coli*, and lambda phage genomes using Bowtie2 in local alignment mode at default settings (*16*) and GraphMap (*17*), respectively. We then ensured only one unique mapping per read, including assigning all lambda reads to the lambda genome (as the complete lambda genome is integrated in the *E. coli* chromosome) prior to calculating relative species proportions.

### *De novo* genome assembly from PacBio and Illumina data

To ensure that our *E. coli* alignments and sequencing measures were not a result of any strain or sample-specific genetic drift or contamination, we performed a *de novo* assembly of the *E. coli* genome used in this study using the PacBio data. We used the Hierarchical Genome Assembly Process (HGAP, v2) for read-cleaning and adapter trimming (pre-assembly), *de novo* assembly with Celera Assembler, and assembly polishing with Quiver (*18*). Raw sequencing reads were filtered for length and quality such that the minimum polymerase read score was 0.8, the minimum subread length was 500 bp, and the minimum polymerase read score was greater than 100. The assembly was generated using CeleraAssembler v1 with the default parameters and was polished using the Quiver algorithm (*18*). Our assembly yielded a single contig of 4,734,145 base pairs at >99.7% accuracy and confirmed the *E. coli* sample as strain K-12.

We next performed independent *de novo* and directed assemblies of the *E. coli* genome using single-end Illumina data. Raw Illumina reads were preprocessed for trimming of adapters and removal of low-complexity and low-quality sequences. *E. coli* reads were identified using Bowtie2 alignment against the *E. coli* reference genome in local alignment mode at default parameters. *De novo* assembly was performed using the SPAdes genome assembly v3.8.2 (*19*) with the “careful” parameter. The *de novo* assembled contigs as well as the original dataset of Illumina reads were then separately mapped to the consensus PacBio *E. coli* genome assembly.

### SURPIrt (sequence-based ultra-rapid pathogen identification, real-time) analysis

The SURPI*rt* pipeline is a real-time analysis pipeline for automated metagenomic pathogen detection and reference-based genomic assembly from nanopore sequencing data. Consisting of an amalgam of previously published SURPI (*6*) and Metapore (*7*) software, and a newly developed SURPI*viz* graphical visualization platform, SURPI*rt* incorporates Linux shell scripts and code from the Python, Perl, Javascript, HTML, and Go programming languages. SURPI*rt* was originally developed for clinical human samples, but was customized for automated analysis of the NASA test mixture of mouse, *E. coli*, and lambda bacteriophage DNA. SURPI*rt* can be run on a server, cloud, or laptop; the data analysis presented here was run a Linux Ubuntu computational server with 64 cores and 1TB memory.

We simulated a real-time SURPI*rt* analysis of the NASA runs using both ground and in-flight nanopore data downloaded from the ISS. In-flight analyses were not possible as a laptop with the necessary computational power and a local basecaller, obviating the requirement for an Internet connection, were not available at the time of these studies. For the simulation, the SURPI*rt* pipeline was run in automated mode to continually scan a download directory containing raw FAST5/HDF files from the MinION sequencer for batch analysis. For a preset number of reads per batch (n=200 for this analysis) corresponding to ~2 minutes of elapsed time, the 2D read or either the template or complement read, depending on which is of higher quality, was extracted from each FAST5 file, and a FASTQ file generated using HDF5 Tools (ref: HDF5/Tools API Specification, http://www.hdfgroup.org/HDF5/doc/RM/Tools.html, accessed September 18th, 2016). Reads corresponding to host mammalian reads (i.e. mouse or human) were subtracted computationally using MegaBLAST (word size 16, e-value cutoff = 10^-5) alignment to the *M. musculus* reference genome. Non-host reads were then aligned to the comprehensive NCBI nt database, which contains GenBank, for identification of viruses, bacteria, fungi, and parasites, respectively. For each read, the single best match by e-value was retained, and the corresponding NCBI Genbank identifier (gi) assigned to the best match was then annotated by taxonomic lookup of the corresponding lineage, family, genus, and species. We tested various configurations of the SURPI*rt* pipeline, including the incorporation of taxonomic classification according to lowest common ancestor (*7*) using either the viral and bacterial portions of the NCBI nt database or the entire NT database.

For automated real-time results visualization (*7*), a live taxonomic count table was generated by SURPI*rt* and displayed as a donut chart using the CanvasJS graphics suite (http://canvasjs.com/, accessed September 26th, 2016) with the chart refreshing every 30 s. For each microbial species detected, the top hit was chosen to be the reference gi in the NCBI nt database (among all reference sequences assigned to that species) with the highest number of aligned reads, with priority given to reference sequences in the following order: (1) complete genomes, (2) complete sequences, or (3) partial sequences or individual genes. Metagenomic heat maps, Krona plots (*8*), coverage maps, and pairwise identity maps were automatically generated using SURPI*rt* for viewing via the SURPI*viz* graphical user interface.

For purposes of comparison, more complete coverage maps were generated by GraphMap alignment to the SURPI*rt*-identified top species gi, and then visualized using SURPI*viz*. In future iterations of SURPI*rt*, a separate computational analysis fork to run GraphMap alignment in real-time will be automatically spawned from species-level “hits” containing a number of reads exceeding a preset threshold.

### *De novo* genome assembly from nanopore data

We used Minimap v0.2-r124-dirty and Miniasm v0.2-r137-dirty as described in the github tutorial, using 8 threads (https://github.com/lh3/miniasm). Two assemblies were generated, using reads that mapped exclusively to *Escherichia coli* K12 MG1655 using GraphMap from the four runs on the ISS, and all reads from the ISS that did not map to host organisms (human and mouse, using genomes hg38 and mm10). We then polished these assemblies using nanopolish v0.5.0. We used Canu v1.73 (*20*) at default parameters using a specified target genome size of 4.8 MB to *de novo* assemble contigs directly from in-flight nanopore data corresponding to the “metagenomic” mixture of mouse, *E. coli*, and lambda genomes. Runs were performed using both a 64-core computational server with 512 gigabytes (Gb) memory and an 8-hyperthreaded core laptop with 64 Gb memory. Four sets of *de novo* assemblies were generated: (1) in-flight runs 1-3 (n=46,871), (2) in-flight runs 1-4 (n=84,502), (3) in-flight runs 1-4 after removal of host *M. musculus* reads (n=53,287), and (4) in-flight runs 1-4 from bacterial reads identified using the SURPI*rt* pipeline. Assembly metrics, including N50, were calculated by the “abyss-stats.pl” program in ABySS (*21*). *E. coli* were mapped to the consensus PacBio genome using Mauve version 2.4.0 (*22*). Genome-wide pairwise identities of *de novo* assemblies were estimated using JSpeciesWS (*23*) after specifying the use of MUMmer (*24*) for pairwise identity calculations.

## Acknowledgements

The Biomolecule Sequencer team is grateful to support personnel at the NASA Johnson Space Center and Marshall Space Flight Center, especially D. Voss and L. Gibson. We would also like to thank the Epigenomics Core Facility of Weill Cornell Medicine and Roger Altman.

## References

1. G. R. Taylor, I. Konstantinova, G. Sonnenfeld, R. Jennings, Changes in the immune system during and after spaceflight. Adv Space Biol Med 6, 1–32 (1997).

2. J. W. Wilson et al., Space flight alters bacterial gene expression and virulence and reveals a role for global regulator Hfq. Proceedings of the National Academy of Sciences of the United States of America 104, 16299–16304 (2007).

3. Y. Feng, Y. Zhang, C. Ying, D. Wang, C. Du, Nanopore-based fourth-generation DNA sequencing technology. Genomics, proteomics & bioinformatics 13, 4–16 (2015).

4. N. J. Loman, J. Quick, J. T. Simpson, A complete bacterial genome assembled de novo using only nanopore sequencing data. Nature methods 12, 733–735 (2015).

5. A. B. R. McIntyre, L. Rizzardi, A. M. Yu, N. Alexander, G. L. Rosen, D. J. Botkin, S. E. Stahl, K. K. John, S. L. Castro-Wallace, K. McGrath, A. S. Burton, A. P. Feinberg, C. E. Mason, Nanopore Sequencing in Microgravity. Nature Partner Journals Microgravity, (In Press 2016).

6. S. N. Naccache et al., A cloud-compatible bioinformatics pipeline for ultrarapid pathogen identification from next-generation sequencing of clinical samples. Genome research 24, 1180–1192 (2014).

7. A. L. Greninger et al., Rapid metagenomic identification of viral pathogens in clinical samples by real-time nanopore sequencing analysis. Genome Med 7, 99 (2015).

8. B. D. Ondov, N. H. Bergman, A. M. Phillippy, Interactive metagenomic visualization in a Web browser. BMC Bioinformatics 12, 385 (2011).

9. H. Li, Minimap: Experimental tool to find approximate mapping positions between long sequences. https://github.com/lh3/minimap/, (2015).

10. S. Goodwin, J. D. McPherson, W. R. McCombie, Coming of age: ten years of next-generation sequencing technologies. Nature reviews. Genetics 17, 333–351 (2016).

11. J. Quick et al., Real-time, portable genome sequencing for Ebola surveillance. Nature 530, 228–232 (2016).

12. A. C. Rand, M. Jain, J. Eizenga, A. Musselman-Brown, H. E. Olsen, M. Akeson, B. Paten, Cytosine Variant Calling with High-throughput Nanopore Sequencing. bioRxiv doi: http://dx.doi.org/10.1101/047134 (2016).

13. J. T. Simpson, R. Workman, P. C. Zuzarte, M. David, L. J. Dursi, W. Timp, Detecting DNA Methylation using the Oxford Nanopore Technologies MinION sequencer. bioRxiv doi: http://dx.doi.org/10.1101/047142 (2016).

14. D. R. Garalde, E. A. Snell, D. Jachimowicz, A. J. Heron, M. Bruce, J. Llyod, A. Warland, M. Pantic, T. Admassu, J. Ciccone, S. Serra, J. Keenan, S. Matrin, L. McNeill, J. Wallace., L. Jayasinghe, C. Wright, J. Blasco, B. Sipos, S. Young, S. Juul, J. Clarke, D. J. Turner in bioRxiv doi: http://dx.doi.org/10.1101/068809 (2016).

15. J. Nivala, D. B. Marks, M. Akeson, Unfoldase-mediated protein translocation through an alpha-hemolysin nanopore. Nature biotechnology 31, 247–250 (2013).

16. B. Langmead, S. L. Salzberg, Fast gapped-read alignment with Bowtie Nat Methods 9, 357–359 (2012).

17. I. Sovic et al., Fast and sensitive mapping of nanopore sequencing reads with GraphMap. Nat Commun 7, 11307 (2016).

18. C. S. Chin et al., Nonhybrid, finished microbial genome assemblies from long-read SMRT sequencing data. Nature methods 10, 563–569 (2013).

19. A. Bankevich et al., SPAdes: a new genome assembly algorithm and its applications to single-cell sequencing. J Comput Biol 19, 455–477 (2012).

20. S. Koren, B. P. Walenz, B. K., J. R. Miller, A. M. Phillippy, Canu: scalable and accurate long-read asembly via adaptive k-mer weighting and repeat separation. bioRxiv, (2016).

21. J. T. Simpson et al., ABySS: a parallel assembler for short read sequence data. Genome Res 19, 1117–1123 (2009).

22. A. E. Darling, B. Mau, N. T. Perna, progressiveMauve: multiple genome alignment with gene gain, loss and rearrangement. PLoS One 5, e11147 (2010).

23. M. Richter, R. Rossello-Mora, F. Oliver Glockner, J. Peplies, JSpeciesWS: a web server for prokaryotic species circumscription based on pairwise genome comparison. Bioinformatics 32, 929–931 (2016).

24. S. Kurtz et al., Versatile and open software for comparing large genomes. Genome Biol 5, R12 (2004).

25. S. F. Altschul, W. Gish, W. Miller, E. W. Myers, D. J. Lipman, Basic local alignment search tool. J Mol Biol 215, 403–410 (1990).

